# Opposing effects of pre-existing antibody and memory T cell help on the dynamics of recall germinal centers

**DOI:** 10.1101/2023.12.15.571936

**Authors:** Ariën Schiepers, Marije F. L. van ’t Wout, Alvaro Hobbs, Luka Mesin, Gabriel D. Victora

## Abstract

Re-exposure to an antigen generates serum antibody responses that greatly exceed in magnitude those elicited by primary antigen encounter, while simultaneously driving the formation of recall germinal centers (GCs). Although recall GCs in mice are composed almost entirely of naïve B cells, recall antibody titers derive overwhelmingly from memory B cells, suggesting a division between cellular and serum compartments. Here, we show that this schism is at least partly explained by a marked decrease in the ability of recall GC B cells to detectably bind antigen. Variant priming and plasmablast ablation experiments show that this decrease is largely due to suppression by pre-existing antibody, whereas hapten-carrier experiments reveal a role for memory T cell help in allowing B cells with undetectable antigen binding to access GCs. We propose a model in which antibody-mediated feedback steers recall GC B cells away from previously targeted epitopes, thus enabling specific targeting of variant epitopes.

## Introduction

Recall antibody responses are critical for resistance to recurring pathogens and essential to the protection afforded by most vaccines,^1–4^ and repeated immunization with the same or variant antigens is a staple of our current approach to vaccination. Recall responses, whose titers often exceed those of primary antibodies by one or two orders of magnitude, are the result of the prolific expansion of prime-derived memory B cells (MBCs) that rapidly differentiate into antibody-secreting plasmablasts (PBs) and plasma cells (PCs) upon re-exposure to an antigen. Several factors contribute to the magnitude and quality of recall antibody responses. In primary germinal centers (GCs), B cells undergo immunoglobulin (Ig) somatic hypermutation and affinity maturation, leading to an overall increase in the affinity of the specific B cell pool.^5^ A fraction of GC B cells is exported into circulation as affinity-matured MBCs, which become the dominant source of the high-affinity antibodies elicited by boosting.^3^ B cells can also be directly exported from primary GCs as high-affinity PB/PCs,^6^ and the antibodies secreted by these cells have been shown to shape subsequent B cell responses.^7^ A number of recent studies show that presence of circulating antibodies to a given epitope prevents B cells with the same specificity from entering GCs and contributing to the secondary antibody pool.^8–12^ Finally, primary responses also produce expanded clones of memory CD4^+^ helper T cells, which are promptly mobilized to provide abundant help to B cells upon boosting.^13^ Recall antibody responses are therefore the product of a complex interplay between the effects of B and T cell memory and those of circulating antibody.

In addition to boosting circulating antibody titers, repeated exposure to antigen also leads to the formation of recall GC reactions. These GCs can in principle be formed either by re-engagement of MBCs, allowing continued evolution of previously matured Igs, or by *de novo* recruitment of naïve B cell clones.^14–16^ Whereas the former would extend affinity maturation and allow for remodeling of B cell clones in response to viral variants, the latter would allow greater *de novo* engagement of naïve B cells, thus countering “original antigenic sin.”^17–19^ Recent studies from our laboratory have identified two prominent but at face value contradictory features of how the mouse immune system navigates this choice.^12,20^ First, we found that, while MBCs are able to re-enter recall GCs as described previously,^21–24^ the large majority of recall GC B cells have no prior GC-experience.^20^ Thus, at the cellular level, secondary GCs appear to favor the *de novo* engagement of naïve B cells. In apparent contrast, our subsequent analysis of antibody titers in serum using a molecular fate-mapping approach showed that recall serum antibodies originate overwhelmingly from MBC clones first expanded during priming, a dominance that persists even after multiple booster doses.^12^ Therefore, at the antibody level, recall responses primarily rely on reutilization of B cell clones originally expanded by the first antigen encounter. Recall immunization thus leads to a schism between a mostly *de novo* GC response a serum antibody response that is almost entirely primary-derived.

Here, we resolve this apparent discrepancy by showing that antibody-mediated feedback suppresses the ability of antigen-binding B cells to enter secondary GCs, resulting in the expansion of a B cell population that fails to detectably bind the immunizing antigen, and thus cannot contribute meaningfully to secondary antibody titers. Secondary GCs fail to form in the absence of antigen-specific memory CD4^+^ T cells, indicating that excess help from memory T cells enables the formation of GCs by normally suboptimal B cell clones. The negative effects of antibody feedback are alleviated by boosting with variant viral antigens, suggesting that pre-existing antibody may promote the focusing of recall GC responses on variant epitopes.

## Results

### Recall GC B cells show impaired binding to antigen

We hypothesized that the failure of recall GCs to contribute to antigen-specific titers upon homologous boosting^12^ might be related to differences in their antigen-binding properties. To investigate this possibility, we followed a strategy we used previously to separate primary and secondary responses by anatomical location.^20^ We first immunized wild-type (WT) C57BL/6 mice or S1pr2-CreERT2.*Rosa26*^Lox-Stop-Lox-tdTomato^ mice (S1pr2-Tomato, which allow us to fate-map primary GC B cells in a tamoxifen-dependent manner^25^) with recombinant hemagglutinin (HA_PR8_) in the right hind footpad (FP) to create a primary GC in the draining popliteal lymph node (pLN). One month later, we boosted these mice in the contralateral (left) FP with the same immunogen to generate a recall response (**Fig. 1A**). We then analyzed the left (boost-draining) pLN by flow cytometry 9 days after HA_PR8_ immunization, when GCs are fully formed in these mice as well as in “primary” control mice primed with the unrelated antigen ovalbumin (OVA) (**Fig. 1B,C**, **Supplementary Fig. 1A**). Whereas a large proportion of B cells in primary GCs bound HA_PR8_ tetramers, binding was almost undetectable in secondary GCs (**Fig. 1D,E**). Fate-mapping of GC B cells in the S1pr2-Tomato mice by tamoxifen treatment during primary GC formation allowed us to measure HA binding separately in secondary GC B cells derived from naïve and memory precursors. Confirming our previous findings,^20^ recall GCs consisted almost entirely of tdTomato^−^ B cells, with only a minor (median 2.1%) contribution from memory-derived tdTomato^+^ cells (**Fig. 1F**). Tetramer binding was negligible also among this small tdTomato^+^ population (**Fig. 1F**, **Supplementary Fig. 1A**).

**Figure 1:**
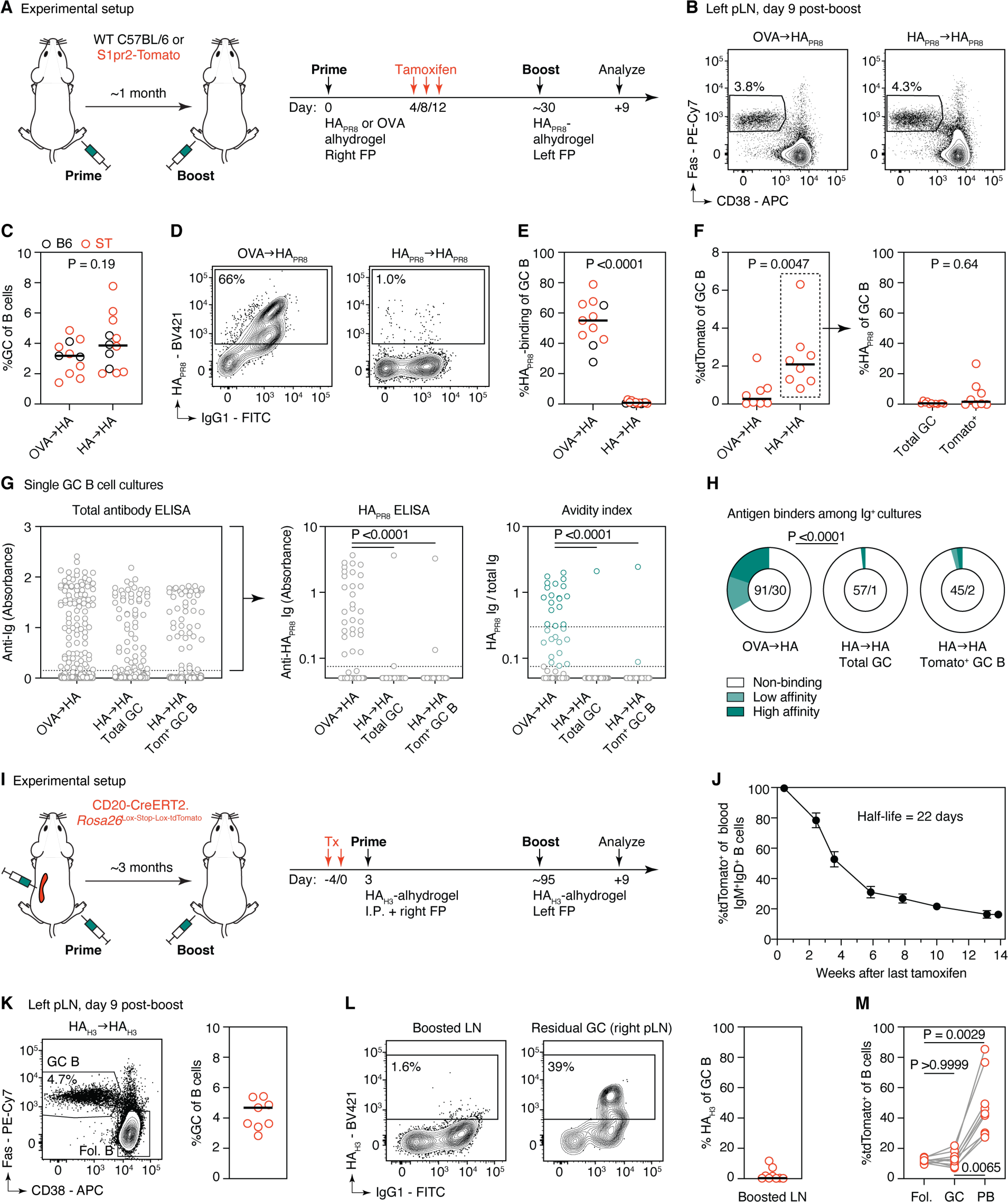
B cells in secondary GCs induced by homologous boosting show impaired binding to antigen. **(A)** Experimental setup for panels (B-H). WT C57BL/6 or S1pr2-Tomato mice were primed in the right FP with HAPR8 or control OVA in alhydrogel adjuvant and boosted ∼1 month later in the left FP with HAPR8-alhydrogel. S1pr2-Tomato mice were given three doses of tamoxifen by oral gavage on the indicated days to label primary GC B cells and their MBC progeny. **(B)** Recall GC formation in the draining pLN, 9 days post-boost. Gated on B cells, as shown in **Supplementary Fig. 1A**. **(C)** Summary of panel B, from 3 independent experiments, with each symbol representing one C57BL/6 (B6, black symbols) or S1pr2-Tomato (ST, red symbols) mouse. n=11 and 12 mice in OVA- and HA-primed groups, respectively. **(D-E)** HAPR8-tetramer binding among GC B cells (D), summarized in (E). **(F)** Memory GC re-entry as defined by percentage of recall GC B cells that are primary-GC derived. **(G)** ELISA measurements of single-GC B cell culture supernatants, as detected by anti-Ig-HRP and coated with either anti-total Ig (left panel) or - HAPR8 (middle panel). The avidity index was calculated by dividing anti-HA by anti-Ig reactivity (right panel), and each Ig^+^ GC B culture was classified as either high-, low- or undetectably binding to HAPR8 as shown. GC B cells were sorted from S1pr2-Tomato mice as detailed in **Supplementary Fig. 1B**. P-values are from Kruskal-Wallis with Dunn’s multiple comparison tests, only significant values (P<0.05) are shown. **(H)** Aggregate data from (G). Numbers are: (Ig+ GC B cell cultures)/(HA binders). P-value is for chi-square test between the indicated groups. **(I)** Experimental setup for panels (J-M). CD20-Tomato mice were given two doses of tamoxifen and primed intraperitoneally and in the right FP three days later with HAH3-alhydrogel. Mice were boosted ∼3 months later in the left FP. Data are from 9 mice from two independent experiments. **(J)** Turnover of the peripheral naïve (IgM+ IgD+) B cell repertoire as assessed over time by flow cytometry of B cells obtained from the blood of CD20-Tomato mice. Gating strategy as shown in **Supplementary Fig. 1C**). Half-life was calculated using one-phase exponential decay analysis. **(K)** Flow cytometry of recall GC B cell formation in the draining pLN, 9 days post-boost. **(L)** HAH3-tetramer binding among GC B cells in the boost-draining pLN as well as a positive-control plot showing a residual GC present in the right pLN (see **Supplementary Fig. 1E**). **(M)** Contribution of tdTomato+ B cells present at the time of priming to follicular (fol.) B cells, GC B cells, and PBs in boosted pLNs. P-values are for Friedman with Dunn’s multiple comparisons test. The solid line in all bar graphs represents the median. P-values in (C, E) and the left panel of (F) are for Mann-Whitney test, and in the right panel of (F) for paired-sample Wilcoxon test.

To rule out that these findings represent an artifact of tetramer staining, we sorted single primary and secondary GC B cells onto tissue culture wells containing feeder cells expressing CD40L, BAFF, and IL-21 (NB-21) that allow single B cells to proliferate and produce assayable amounts of antibody^26^ (**Supplementary Fig. 1B**). We then assayed wells producing detectable amounts of Ig for their ability to bind HA_PR8_ by ELISA (**Fig. 1G,H**). To estimate binding affinity, we calculated the ratio of HA_PR8_ binding to Ig concentration, or “avidity index,”^26^ categorizing samples into 3 strata (below detection, low, and high binding). Although the majority of supernatants from all experimental groups did not react detectably with the immunizing antigen in this assay, reactivity was much lower among GC B cell supernatants from secondary GCs compared to primary GCs, which was true for both total GC and tdTomato^+^ sorted B cells (**Fig. 1G,H**). Thus, primary exposure to an antigen skews the binding properties of B cells recruited to secondary GCs, leading to loss of positivity in both tetramer binding and ELISA assays.

### Failure of recall GC B cells to detectably bind antigen is not due to depletion of the naïve repertoire

A potential explanation for the skewed binding pattern of B cells in secondary GCs is that primary immunization may have depleted the naïve repertoire of B cells specific for the immunizing antigen. To rule this out, we allowed the naïve B cell pool time to re-form prior to boosting, using CD20-CreERT2.*Rosa26*^Lox-Stop-Lox-tdTomato^ (CD20-Tomato) mice^27^ to measure naïve B cell turnover. We first fate-mapped the entire mature B cell compartment in these mice with two consecutive doses of tamoxifen (**Fig. 1I**). Three days later, when virtually all circulating IgM^+^IgD^+^ naïve B cells expressed tdTomato (**Fig. 1J**, **Supplementary Fig. 1C**), mice were immunized with HA_H3_ both intraperitoneally and in one footpad (**Fig. 1I**). The fraction of fate-mapped naïve B cells declined progressively thereafter, with a median of 51.1% and 30.9% cells labeled after 25 and 41 days respectively, yielding a calculated half-life of 21.7 days (**Fig. 1J**). Turnover slowed down after 6 weeks—possibly reflecting the existence of a long-lived IgM^+^IgD^+^ population—stabilizing at approximately 16.0% at 14 weeks post-tamoxifen, including a minor contribution of spontaneous *Rosa26*^Lox-Stop-Lox-tdTomato^ recombination in the absence of tamoxifen estimated at 1.6% (**Supplementary Fig. 1D**). Boosting fate-mapped mice on the contralateral footpad at 3 months post-priming still generated GCs in which very few B cells (median 0.4%) were able to bind the HA_H3_ tetramer (in contrast to occasional residual primary GCs with high binding) (**Fig. 1K,L, Supplementary Fig. 1E**). Therefore, the failure of secondary GCs to detectably bind the immunizing antigen is not due to depletion of antigen-binding precursors from the naïve pool by primary immunization. Of note, these experiments also strengthen our previous finding that secondary GCs are composed almost exclusively of naïve-rather than memory-derived B cells:^20^ whereas recall PBs were markedly enriched for tdTomato^+^ cells compared to follicular B cells in the same LN, indicative of their memory origin, the same was not true for secondary GC B cells, which were fate-mapped to the same extent as naïve B cells in the same mice (**Fig. 1M**). Thus, recall GCs are predominantly composed of B cells that had not yet been generated at the time of priming and are therefore unequivocally naïve-rather than memory-derived.

### Failure of recall GC B cells to bind antigen is caused by pre-existing antibody

Serum antibodies have been shown to compete at the epitope level with naïve B cells of the same fine specificity, preventing their entry into GCs.^7–10,23^ We hypothesized that, if loss of antigen binding in recall GCs is related to this phenomenon, mice boosted heterologously with antigenically drifted HAs containing novel epitopes not targeted by primary antibody would generate secondary GCs with improved binding to antigen. To test this, we primed mice with HA_FM1_, an HA variant with 10% amino acid divergence from HA_PR8_, and boosted these mice with HA_PR8_ as above (**Fig. 2A**). HA_PR8_-binding was partially rescued in GCs of heterologously boosted compared to homologously boosted mice, with no change in the ability of MBCs to enter secondary GCs (**Fig. 2B-D**). These results were corroborated by ELISA measurements on the supernatants of single-GC B cell cultures (**Fig. 2E,F**). Rescue of HA_PR8_-binders was greater when mice were primed with HA_CA’09_ and boosted with HA_PR8_ (representing a 20% amino acid-level divergence), consistent with a dose-dependent effect of antigenic distance (**Fig. 2G-I**).

**Figure 2:**
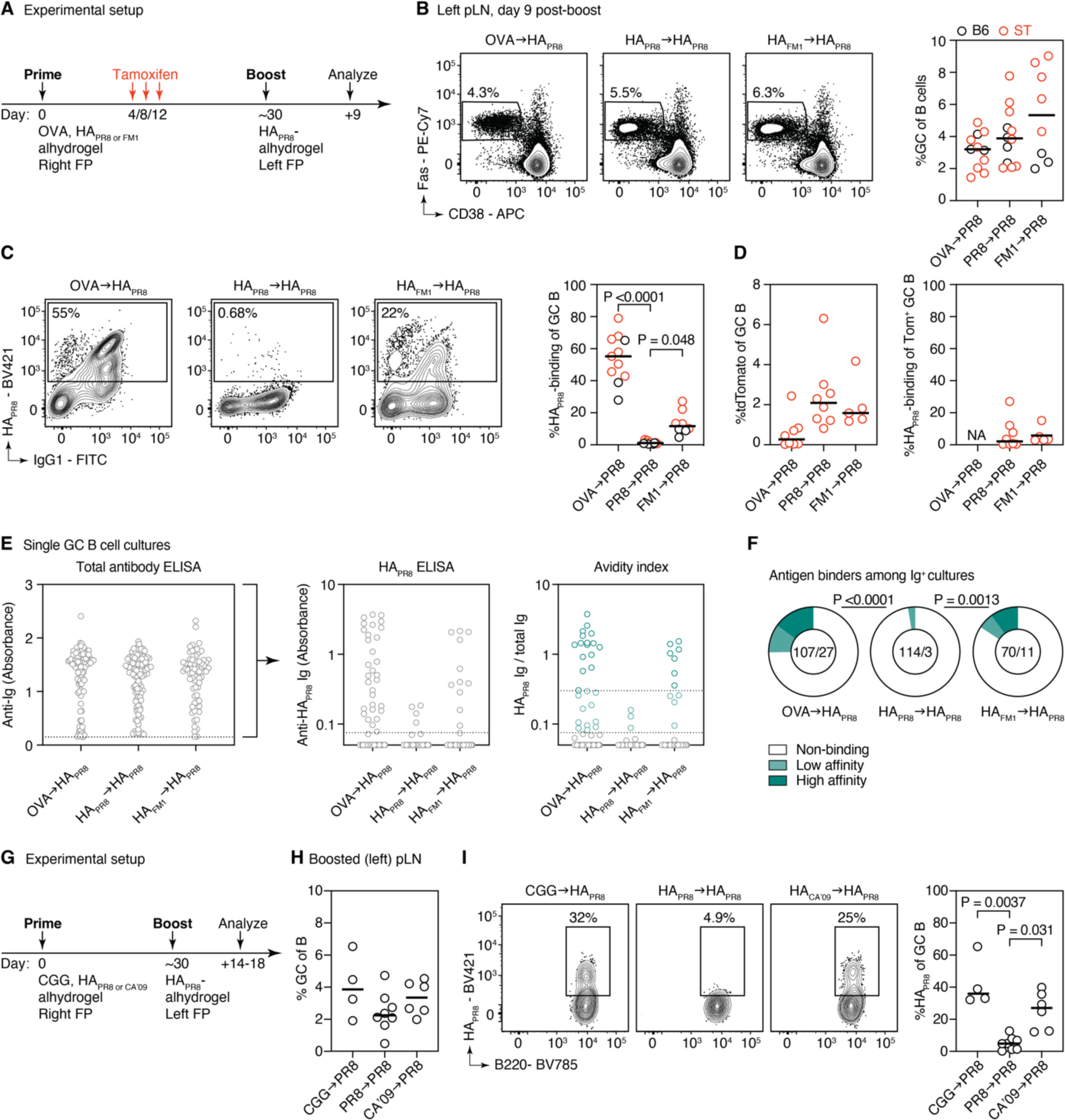
Rescue of recall GC B cell antigen binding upon boosting with variant HAs. **(A)** Experimental setup as in Fig. 1A, with an additional third group of mice primed with HAFM1-alhydrogel (8 mice from 2 independent experiments). This experiment was conducted in parallel with those shown in Fig. 1. OVA→ HAPR8 and HAPR8→ HAPR8 data are reproduced from Fig. 1. **(B-D)** Recall GC formation (B), HAPR8-tetramer binding (C), and GC re-entry of tdTomato^+^ MBCs (D) in the draining pLN, 9 days post-HAPR8 boost. **(E)** ELISA of single-GC B cell culture supernatants sorted from C57BL/6 mice. Only Ig^+^ supernatants are shown. **(F)** Aggregate data from (E). Numbers are: (Ig^+^ GC B cell cultures)/(HA binders). **(G)** Experimental setup for (H, I). Mice were primed with HACA’09, HAPR8, or an irrelevant control (chicken gamma globulin, CGG) and boosted HAPR8 in the contralateral FP ∼1 month later. Data are from 4-8 mice per group from at least two independent experiments. **(H, I)** Flow cytometry for recall GC formation (H) and HAPR8-tetramer binding among GC B cells (I). The solid line in all bar graphs represents the median. P-values are for chi-square tests between indicated groups (F) and for significant (p<0.05) Kruskal-Wallis with Dunn’s multiple comparisons (C, I).

To formally test whether these differences were due to antibody-mediated feedback, we used *Aicda*^CreERT2/+^*.Rosa26*^Confetti/Confetti^.*Prdm1*^flox/flox^ mice (*Prdm1*ΔGC), in which the gene required for PB/PC differentiation (*Prdm1*, encoding for the transcription factor Blimp-1) can be deleted in a tamoxifen-inducible manner from the cohort of B cells that responded to priming, rendering these cells and their descendants incapable of differentiating into antibody-secreting cells. We primed and boosted *Prdm1*ΔGC or control *Aicda*^CreERT2/+^*.Rosa26*^Confetti/Confetti^.*Prdm1*^+/+^ mice (AID-Confetti^28^) with HA_PR8_ in alhydrogel, as above (**Fig. 3A**). Anti-HA_PR8_ IgG was reduced by approximately 9.0 and 5.4-fold in *Prdm1*^flox/flox^ compared to *Prdm1*^+/+^ mice prior to and after boosting, respectively (**Fig. 3B**). This partial depletion of antigen-specific antibodies resulted in a partial rescue of the secondary GC phenotype, with secondary GCs in *Prdm1*ΔGC mice containing a median of 14.3% HA-binding B cells compared 1.5% in control mice (**Fig. 3C,D**). HA-binding among secondary GC B cells showed a moderate but significant inverse correlation with anti-HA IgG titers in homologously-boosted *Prdm1*ΔGC mice (r^2^=0.35, P=0.035, **Supplementary Fig. 2A)**. Thus, primary-derived antibodies directly inhibit the participation of high-affinity B cells in recall GCs.

**Figure 3:**
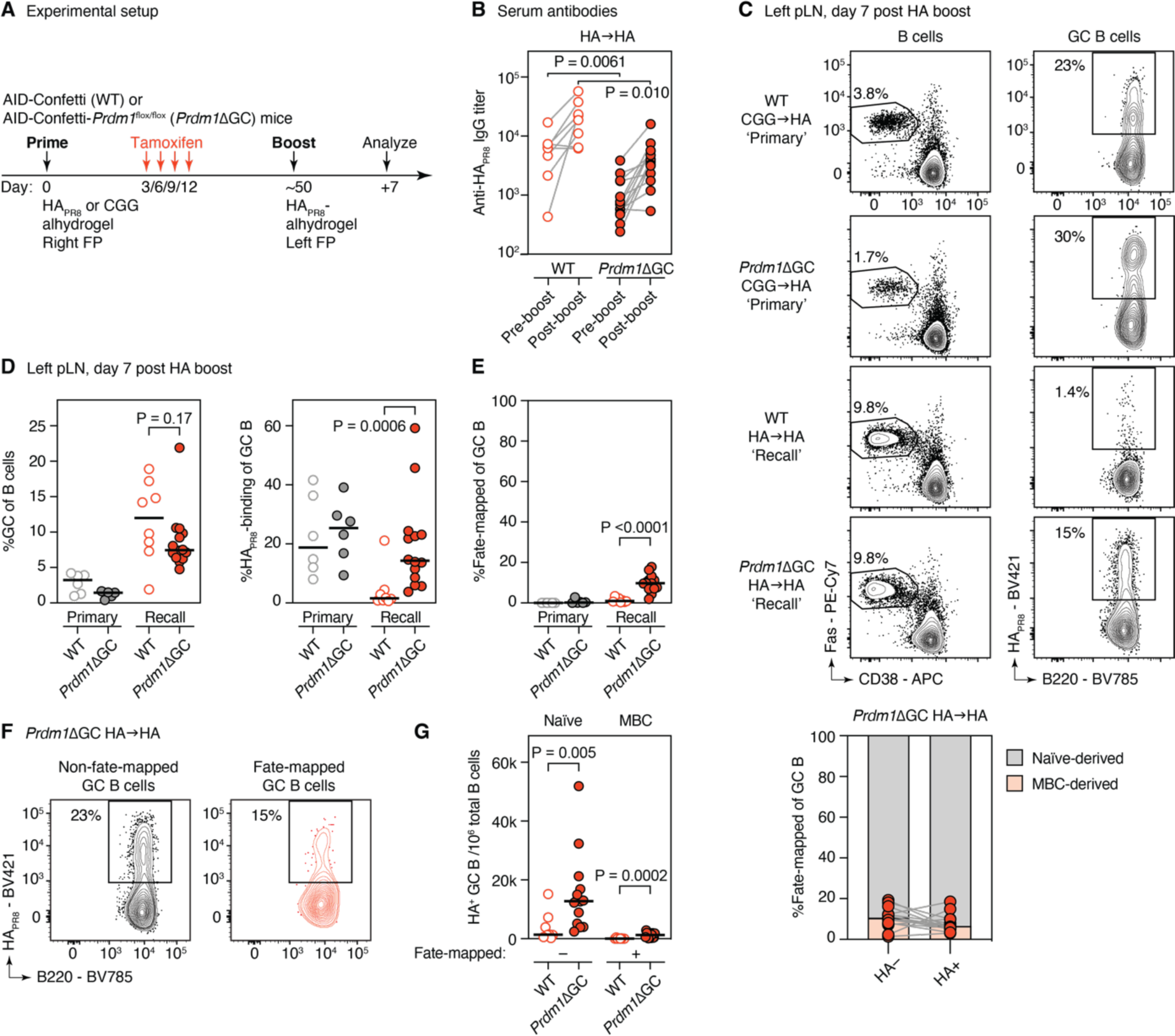
Depletion of primary antibodies increases antigen-binding in secondary GCs. **(A)** Experimental setup. AID-Confetti mice either *Prdm1*^flox/flox^ (*Prdm1*ΔGC) or *Prdm1*-sufficient (“WT”) were primed in the right FP with HAPR8 or irrelevant CGG-alhydrogel, given 4 doses of tamoxifen by oral gavage between days 3 and 12 to fate-map the progeny of activated (AID-expressing) B cells and ablate *Prdm1* in *Prdm1*ΔGC mice. All mice were boosted in the contralateral FP with HAPR8-alhdydrogel. Data are from 6-14 mice from two independent experiments. **(B)** Anti-HAPR8 IgG endpoint titers before and 1 week after boosting with HAPR8-alhydrogel as determined by ELISA. **(C)** Representative plots of GC formation (left panel, pre-gated on B cells) and HA-tetramer binding among GC B cells (right panel), in draining (left) pLN 1 week post-boost. **(D, E)** Quantification of data as in (C). **(F)** Representative plots showing HA binding among fate-mapped and non-fate-mapped GC B cells in *Prdm1*ΔGC mice. **(G)** Quantification of (F). The solid lines represent the median. P-values comparing *Prdm1*ΔGC and *Prdm1*-sufficient mice are for Kruskal-Wallis with Dunn’s multiple comparisons tests at pre- and post-boost timepoints (B) and Mann-Whitney tests (D, E, G).

The presence of a fate-mapping *Rosa26*^Confetti/Confetti^ allele in these mice also allowed us to assess the re-entry of MBCs into secondary GCs. The fraction of memory-derived B cells was significantly higher in *Prdm1*ΔGC mice (median 9.8%) compared to controls (median 0.93%) (**Fig. 3D**). Because Blimp-1-deficient B cells are precluded from differentiating into PB/PCs, loss of this factor may affect both MBC export from the primary GC and the decision of MBCs to re-enter GCs. These experiments therefore do not allow us to determine the extent to which antibody feedback is responsible for the increase in MBC-derived cells in secondary GCs. Nevertheless, the large majority (>90%) of recall GC B cells remained non-fate-mapped—and are therefore of naïve origin—even when circulating antibody is depleted (**Fig. 3E**). Importantly, an equally high proportion of HA^+^ GC B cells was non-fate-mapped in *Prdm1*ΔGC mice, and the increase in HA^+^ GC B cells was mostly observed in non-fate-mapped cells, indicating that the rescue in detectable binding was mostly due to enhanced recruitment of HA-binding naïve cells rather than MBCs (**Fig. 3F,G**). We conclude that, while antibody-mediated feedback cannot explain the dominance of naïve-derived B cells in secondary GCs, it does account at least partially for the loss of antigen-binding among this population.

### Memory T cells are required for recall GCs to form in the presence pre-existing antibody

A potential contributor to the ability of B cells with poor antigen binding capacity to enter recall GCs is the presence of a pre-expanded population of memory T cells, which would act by lowering the affinity threshold required for GC entry.^29,30^ To test this possibility, we employed a hapten-carrier model where T cell help can be de-coupled from B cell specificity. We induced GCs by footpad immunization with 4-hydroxy-3-nitrophenylacetyl (NP)-CGG in alhydrogel of WT mice previously primed in the opposite leg with OVA (*primary* group), NP-CGG (*homologous recall* group), or NP-OVA (*carrier-switch* group) (**Fig. 4A**). In both recall groups, antibodies and MBCs to the hapten NP will be present at the time of boosting, whereas memory T cells to the carrier protein CGG will only be present in the homologous recall group. As with protein immunization, NP-CGG induced abundant GCs in the homologous recall group, which displayed moderately but significantly reduced tetramer binding compared to primary GCs (**Fig. 4B,C**). Usage of Igλ^+^ B cells, characteristic of an anti-NP response, was also significantly reduced in recall GCs, but remained enriched compared to the available B cell repertoire (which is ∼5% Igλ^+^ in C57BL/6 mice^31^) (**Supplementary Fig. 2B**). More dramatically, formation of secondary GCs in the carrier-switch group was completely abrogated (**Fig. 4B,C**), indicating that help from naïve CGG-specific T cells is insufficient to overcome the negative effect of anti-NP antibody on GC formation. Moreover, PB formation was markedly inhibited in carrier-switch mice compared to the homologous group (**Fig. 4B,C**), indicating that T cell help is required for the MBC to PB transition in this setting. To test whether the inability of carrier-switch mice to generate recall GC and PB responses could be directly attributed to the presence of anti-NP antibodies, we immunized *Prdm1*ΔGC and control AID-Confetti mice with OVA (primary group) or NP-OVA (carrier-switch group) and administered tamoxifen during the primary response to prevent PB/PC formation (**Fig. 4D**). NP-specific IgG titers were strongly if not fully depleted in *Prdm1*ΔGC mice compared to controls at both pre- and post-boost timepoints (6.3- and 7.3-fold reduction, respectively; **Fig. 4E**). In carrier-switch settings, boosting with NP-CGG resulted in significantly increased GC B cell and PB populations in *Prdm1*ΔGC compared to *Prdm1*-sufficient mice (**Fig. 4F**), with anti-NP IgG titers inversely correlating with GC size (r^2^=0.43, P=0.0032, **Supplementary Fig. 2C**), indicating that a partial reduction in anti-NP antibodies partially rescues GC formation also in the absence of T cell memory (**Fig. 4F**). In this setting, depletion of antibodies led to no increase in the participation of MBCs in recall GCs (**Fig. 4G**).

**Figure 4:**
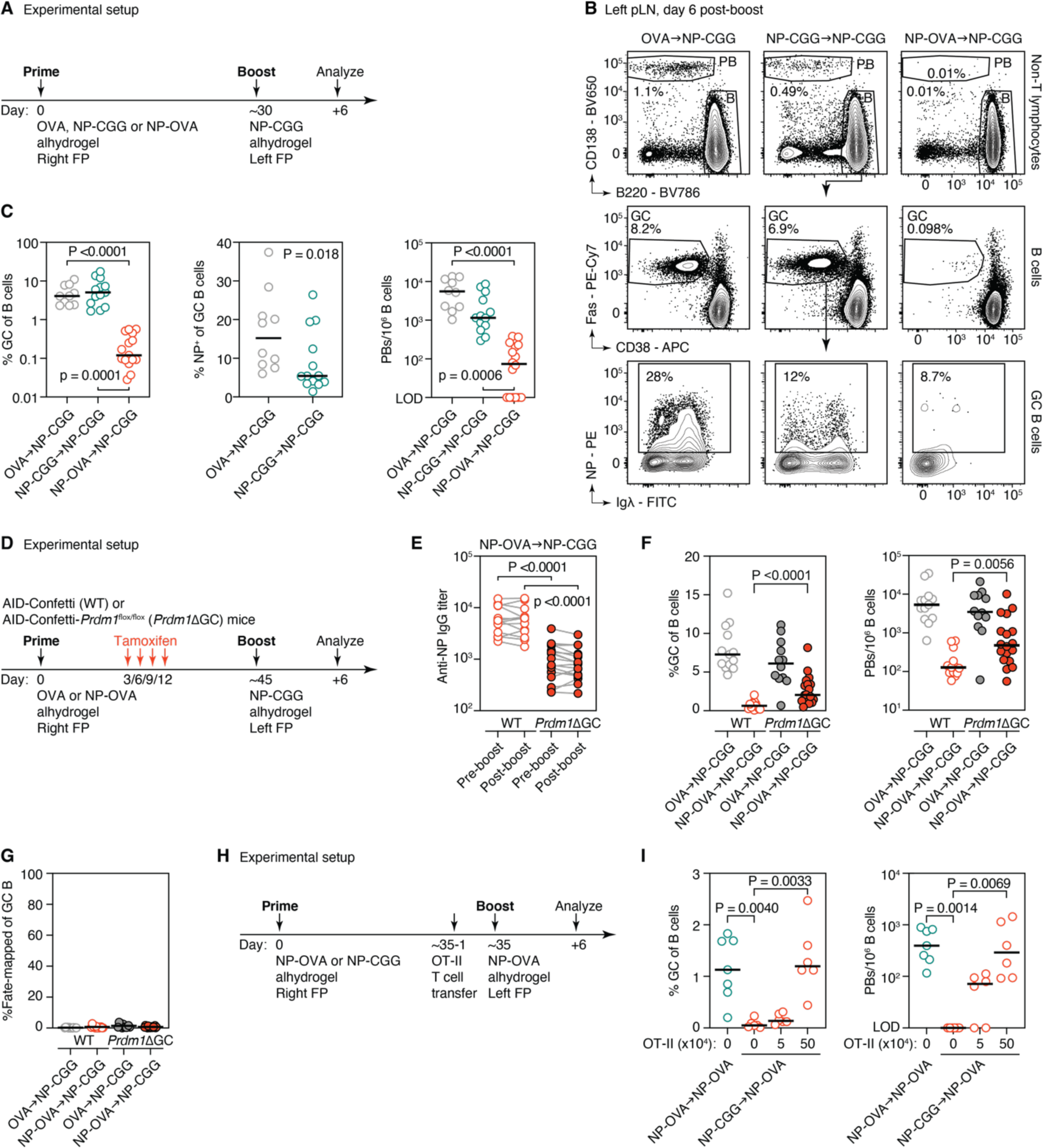
Memory T cells are required for secondary GC B cell formation in the presence of antibody feedback. **(A)** Experimental setup for panels (B,C). C57BL/6 mice were boosted with NP-CGG in alhydrogel ∼1 month after immunization in the contralateral FP with either OVA (primary group), NP-CGG (homologous recall group), or NP-OVA (carrier-switch group) adjuvanted with alhydrogel. Data are from 10-15 mice per group from at least three independent experiments. **(B-C)** Flow cytometry of lymphocytes in draining left pLN 6 days after boost. (B) Representative plots of PBs and B cells, pre-gated on non-T cells (top row), GC formation (middle row) and NP-binding of GC B cells (bottom row) are shown, quantified in (C). **(D)** Experimental setup for panels (E-G). AID-Confetti mice either *Prdm1*^flox/flox^ (*Prdm1*ΔGC) or *Prdm1*-sufficient (“WT”) were primed in the right FP with either OVA (primary group) or NP-OVA in alhydrogel (carrier switch group) and administered tamoxifen 4 times between days 3 and 12 prior to boosting at day ∼45 day in the left FP with NP-CGG in alhydrogel. Data are from 12-18 mice per group from three independent experiments. **(E)** Anti-NP IgG endpoint titers before and 6 days after boosting as determined by ELISA. **(F-G)** Flow cytometry of draining pLN 6 days post-boost. GC formation (F, left panel), PB induction (F, right panel) and MBC re-entry into recall GCs (G) were quantified. **(H)** Experimental setup for (I). Similar as in A-C, C57BL/6 mice were boosted with NP-OVA in alhydrogel ∼1 month after immunization in the contralateral FP with either NP-OVA (homologous recall group) or NP-CGG (carrier-switch group) adjuvanted with alhydrogel. One day before boosting, 0, 5 or 50 *10^4^ T cells purified from monoclonal OT-II mice were transfer intravenously into carrier-switch group mice. Data are from 6-7 mice per group and from two independent experiments. **(I)** Flow cytometric analysis of GC (left) and PB formation (right) 6 days after boosting with NP-OVA. The solid line in all bar graphs represents the median. The P-values represent significant (p<0.05) Kruskal-Wallis with Dunn’s multiple comparisons (C, left and right panel; E; and I) and Mann-Whitney tests (C, middle panel; and F, where *Prdm1*ΔGC and *Prdm1*-sufficient mice are compared).

Finally, we sought to confirm that the ability of secondary GCs to form in the face of antibody feedback is a consequence of the increased availability of helper T cells. To this end, we used adoptive transfer of monoclonal T cells to increase helper T cell availability prior to boosting in carrier-switch settings, where GCs fail to form (**Fig. 4H**). We primed mice with NP-CGG (carrier-switch) or NP-OVA (homologous recall), which we boosted one month later in the contralateral footpad with NP-OVA. Carrier-switch mice received either no adoptive transfer or a low (5 x 10^4^) or high (5 x 10^5^) number of congenically-marked (CD45.1) OVA-specific OT-II T cells one day prior to boosting (**Fig. 4H**, **Supplementary Fig. 2D,E**). Whereas formation of recall GCs and PBs was again strongly suppressed in carrier-switch mice that did not receive OT-II cells, this was overcome by adoptive transfer of high, but not low, numbers of OT-II T cells (**Fig. 4I**), with the abundance of transfer-derived T cells correlating positively with GC size (r^2^=0.83, P<0.0001, **Supplementary Fig. 2F**). NP-binding remained low in all recall settings (**Supplementary Fig. 2G**). Thus, increasing T cell help overcomes the suppressive effect of antibody feedback on GC formation, but not on high-affinity B cell recruitment. We conclude that, in homologous boosting regimens, the increased availability of help from memory T cells allows GCs to form even in the presence of antibody-mediated feedback, but at the cost of a loss of effective antigen binding among recruited B cells.

## Discussion

Collectively, our experiments support a model in which the binding properties of B cells in recall GCs are shaped by the opposing forces of antibody-mediated feedback and T helper cell memory. In agreement with prior studies,^7–10,23^ serum antibody exerts a suppressive effect by blocking access to GCs by naïve B cells with the same specificity. In the absence of pre-expanded memory T cell clones, as achieved experimentally in our carrier-switch experiments, suppression by circulating antibody does not allow GCs to form. However, in the presence of memory T cell help, GCs form efficiently but are populated by B cells that bind antigen poorly if at all. We speculate that poor antigen binding among secondary GC B cells may in part be due to antibody-mediated masking of immunodominant epitopes directing these cells towards cryptic/non-immunodominant sites^26^ or degradation products,^32^ making them difficult to detect by tetramer staining. Alternatively, but not exclusively, excess T cell help may allow B cells with very low affinity for antigen (or even no affinity at all^30^) to efficiently access secondary GCs. Access of low-affinity B cells can be further increased by formation of multivalent complexes between antigen and pre-existing antibody.^10^ Our finding that secondary GCs elicited by NP boosting are enriched in B cells that are most likely NP-specific (given that they are Igα^+^) yet fail to bind NP by flow cytometry is in line with this possibility. Low-affinity or otherwise altered antigen binding may also underlie the failure of secondary GC B cells to produce detectable antibody responses in serum,^12^ either because these cells are not of sufficient quality to allow substantial PB/PC differentiation or because the antibody their progeny secrete is not of high enough affinity to be detected by conventional methods.

In contrast to homologous boosting, *de novo* B cell responses induced by variant antigens specifically target escape epitopes not masked by circulating antibody.^12^ Our findings suggest that antibody-mediated suppression of crossreactive responses serves the critical function of specifically guiding *de novo* GCs towards variant-specific epitopes, thus weakening imprinting effects such as original antigenic sin.^17,18^ Within this framework, the fundamental role of recall GCs, enforced by antibody-mediated feedback, would be to mature naïve-derived B cells specifically tailored to escape epitopes on viruses or other pathogens, rather than to allow secondary maturation of the memory repertoire. Our antibody depletion experiments indicate this would be achieved mainly by antibody-mediated suppression of naïve B cells with overlapping specificities, rather than by specifically blocking GC re-entry by MBCs. Along these same lines, the GCs with poor antigen binding that arise upon repeated boosting with the same antigen would represent aberrant byproducts of a system geared towards responding to heterologous rather than homologous challenge.

Lastly, our experiments using CD20-Tomato mice rule out the possibility that our prior finding of inefficient recruitment of fate-mapped B cells to secondary GCs^20^ was due to failure of the S1pr2-CreERT2 or *Aicda*^CreERT2^ drivers to fate-map early, pre-GC MBCs. Because we could not rule this out in our previous study, we referred to the B cell precursors of secondary GCs as “likely naïve.”^20^ In our CD20-Tomato experiments, most B cell clones that populate secondary GCs had not even emerged from the bone marrow by the time of priming. These findings confirm that, at least in mice, the dominant B cell population in boost-induced recall GCs is indeed the progeny of truly naïve precursors.

## Acknowledgments

We thank all members of the Victora Laboratory for helpful discussion, L. Shen and J. Pae for technical assistance, C.L. Ferreira and J. Bortolatto for mouse colony management, the Rockefeller University Comparative Biosciences Center for mouse housing, and all Rockefeller University staff for their continuous support. We thank K. Gordon and J.-P. Truman for single-cell sorting, G. Kelsoe (Duke University) for NB-21.2D9 cells, T. Kurosaki and T. Okada (Osaka University and RIKEN-Yokohama) for S1pr2-CreERT2 mice, M. Shlomchik (U. Pittsburgh) for CD20-CreERT2 mice, and M. Nussenzweig, A. Barbulescu and S.T. Chen for critical reading of the manuscript. This study was funded by NIH/NIAID grants R01AI119006, R01AI157137, and R01AI173086 to G.D.V. Work in the Victora laboratory is additionally supported by NIH grant DP1AI144248 (Pioneer award), the Robertson Foundation, and the Stavros Niarchos Foundation Institute at Rockefeller University (SNIFRU). A.S. was supported by a Boehringer-Ingelheim Fonds PhD fellowship. G.D.V. is a Burroughs-Wellcome Investigator in the Pathogenesis of Infectious Disease.

## Author Contributions

A.S. performed most experimental work and data analysis, with help from M.F.L.v.’t.W. and essential input from L.M.. Nojima culture experiments were performed and analyzed by M.F.L.v.’t.W. and A.S. with assistance from A.H.. A.S., L.M. and G.D.V. conceptualized the study. A.S. and G.D.V designed all experiments, interpreted the data and wrote the manuscript. All authors reviewed and edited the final manuscript.

## Methods

### Mice

Wild-type C57BL/6J (CD45.2), *Rosa26*^Lox-Stop-Lox-tdTomato^ (AI14^33^) and *Rosa26*^Confetti 34^ mice were purchased from The Jackson Laboratory (strain numbers: 000664, 007914 and 013731, respectively). Strains that were kindly provided to us: *Aicda*^CreERT2 21^ by J.-C. Weill and C.-A. Reynaud (Université Paris-Descartes), *S1pr2*^CreERT2^ BAC-transgenic ^25^ by T. Kurosaki and T. Okada (U. Osaka, RIKEN-Yokohama), CD20-CreERT2^27^ by M. Shlomchik (U. Pittsburgh) and *Prdm1*^flox/flox 35^ by J. Boss (Emory University). OT-II TCR transgenic (Y-chromosome)^36^ CD45.1 mice were bred and maintained in our laboratory. All mice were held at the immunocore clean facility at the Rockefeller University under specific pathogen-free conditions. All mouse procedures were approved by the Rockefeller University’s Institutional Animal Care and Use Committee.

### Immunizations and treatments

Immune responses were induced in 6-12-week-old male and female mice by subcutaneous immunization in the right FP with 5 µg (for HA experiments) or 10 µg (for hapten-carrier experiments) supplemented with 1/3 volume alhydrogel adjuvant (Invivogen). In S1pr2-Tomato mice, the primary GC B cell response was fate-mapped by oral gavage of 200 µl tamoxifen (Sigma) dissolved in corn oil at 50 mg/ml, on days 4, 8 and 12. For the CD20-Tomato experiment, mice were given two doses of 250 µl tamoxifen prior to FP (5 µg) as well as intraperitoneal (20 µg) immunization with HA_H3_-alhydrogel. For Blimp-1-depletion experiments, 200 µl tamoxifen was given on days 3, 6, 9, and 12. Blood samples were collected in these mice pre and post-boost, via cheek puncture into microtubes prepared with clotting activator serum gel (Sarstedt). To induce recall B cell responses, mice were boosted in the contralateral (left) FP at the timepoints detailed in the figures and figure legends. For adoptive T cell transfer, spleens of naïve CD45.1 OT-II mice were harvested and homogenized by filtering through a 70-μm cell strainer and red-blood cells were lysed with ACK buffer (Thermo Scientific). CD4^+^ T cells were isolated by negative selection using a cocktail of biotinylated antibodies targeting Ter119, CD11c, CD11b, CD25, B220, NK1.1, and CD8, followed by anti-biotin beads (Miltenyi Biotec), as per the manufacturer’s instructions. Subsequently, isolated CD4^+^ T cells were injected intravenously in 100 μl PBS per mouse.

### Generation of recombinant proteins

Recombinant HAs used for immunizations were produced in-house using a CHO cell protein expression system, as described previously.^20^ Cysteine residues were introduced into the HA sequence to create trimer-stabilizing disulfide bonds, as originally described by Ian Wilson’s lab.^37^ We produced HA_PR8_, HA_CA’09_ and HA_FM1_ previously,^12,20^ and for HA_H3_ (H3/A/Wisconsin/67/2005) the same procedure was followed, including the introduction of trimer-stabilizing mutations. For immunizations, C-terminal domains not native to HA (foldon, Avi-tag, His-tag) were removed by thrombin cleavage and HAs were subsequently FPLC-purified prior to storage in phosphate-buffered saline (PBS). For ELISA, non-thrombin treated FPLC-purified protein was used. HA_PR8_ tetramers for flow cytometry were generated by site-specific biotinylation of non-cysteine-stabilized treated HA protein containing the Y98F mutation that prevents sialic acid binding using BirA-500 ligase (Avidity), followed by Zeba desalting column purification (Thermo Fisher). Biotinylated HA was incubated with Streptavidin-BV421 in PBS for 30 min at room temperature at a molar ratio of 4 to 1 (HA-trimer to Streptavidin). The plasmid used for HA cloning and expression (pVRC8400) and HA_H3_ protein for tetramer construction were kindly provided by A. McDermott (VRC/NIAID/NIH). All other proteins were obtained commercially: NP-CGG, NP-OVA, OVA (Biosearch), CGG (Rockland Immunochemicals).

### Flow cytometry and sorting

For flow cytometry and cell sorting, lymph node cell suspensions were obtained by mechanical disassociation with disposable micropestles (Axygen). Cells were resuspended in PBS supplemented with 0.5% BSA and 1 mM EDTA and incubated first with Fc-block (rat anti-mouse CD16/32, clone 2.4G2, Bio X Cell) for 30 min on ice and subsequently with various fluorescently-labeled antibodies (see **Supplementary Table 1**) for 30-60 min. Cells were filtered and washed with the same buffer before analysis on a BD FACS Symphony A5 cytometer or single-cell sorted using a BD FACS Symphony S6. Data were analyzed using FlowJo v.10 software.

### Single GC B cell cultures

NB-21.2D9 feeder cells expressing CD40L, BAFF, and IL-21 (kindly provided by G. Kelsoe, Duke University) were cultured in DMEM supplemented with 10% heat-inactivated FBS and penicillin streptomycin solution (Corning). One day before single-cell sorting, the cells were detached and resuspended in OptiMEM, irradiated (20 Gy) and seeded into 96-well plates at 3,000 cells per well in OptiMEM supplemented with 10% heat-inactivated FBS, 2 mM L-glutamine, 1 mM sodium pyruvate, 50 μM 2-ME, penicillin streptomycin solution, 10 mM HEPES, MEM vitamin solution (Sigma) and MEM non-essential amino acids (Gibco). The following day, single GC B cells were sorted into wells and 150 μl of supplemented OptiMEM along with 30 μg/ml LPS (Sigma-Aldrich, #L6511) and 4 ng/ml IL-4 (Fisher Scientific, #404-ML) was added to each well. Supernatants were harvested 7 days after sorting and screened for Ig and HA_PR8_ reactivity by ELISA, as described below.

### ELISA

To determine antibody levels in supernatants of single GC B cell cultures or in the serum of immunized mice, ELISAs were performed as described before.^20^ 96-well high-binding half-area microplates (Greiner or Corning) were coated overnight at 4°C with antigen or capture antibody in PBS (25 μl per well). In between each step, plates were washed with PBS + 0.05% Tween20. After overnight incubation, plates were blocked with 2.5% bovine serum albumin (BSA, Sigma) in PBS for 2 hours at room temperature. For single GC B cell cultures experiments, 25 μl of undiluted supernatant was added to wells coated with 1 μg/ml goat anti-mouse Ig (Southern Biotech) or HA_PR8_ and for serum samples 3-fold serial dilutions of serum samples starting at 1/100 were added to wells coated with NP_4_-BSA (10 μg/ml, Biosearch) or HA_PR8_ (1 μg/ml). After washing, detection occurred with goat anti-mouse Ig-HRP (Southern Biotech) or IgG-HRP (Jackson Immunoresearch) for single cell cultures and serum ELISAs respectively, followed by development with TMB (slow kinetic form, Sigma). The reaction was stopped with 1N hydrochloric acid and the absorbance was measured at 450 nm on a Fisher Scientific accuSkan FC plate reader.

**Supplementary Table 1:** Flow cytometry antibodies.

| Marker | Fluorophore | Clone | Vendor | Catalog | Dilution |
| --- | --- | --- | --- | --- | --- |
| B220 | BV785 | RA3-6B2 | BioLegend | #103246 | 1:400 |
| TCRβ | APC/eFluor780 | H57-697 | eBioscience | #47-5961-82 | 1:400 |
| CD138 | BV650 | 281-2 | BioLegend | #142517 | 1:400 |
| CD38 | APC | 90 | eBioscience | #17-0381-82 | 1:400 |
| FAS/CD95 | PE-Cy7 | Jo2 | BD Biosciences | #557653 | 1:800 |
| GL7 | FITC | GL7 | BD Biosciences | #553666 | 1:200 |
| CD4 | BV500 | RM4-5 | BD Biosciences | #560782 | 1:200 |
| CD8a | BV500 | 53-6.7 | BD Biosciences | #560776 | 1:200 |
| PD1 | APC-Cy7 | 29F.1A12 | BioLegend | #135224 | 1:200 |
| CXCR5 | BV650 | L138D7 | BioLegend | #145517 | 1:100 |
| NP | PE | N/A | Biosearch Technologies | #N-5070 | 1:200 |
| CD45.1 | BV421 | A20 | BioLegend | #110731 | 1:200 |
| CD45.2 | APC | 104 | BioLegend | #109814 | 1:200 |
| IgD | BV650 | 11-26c.2a | BioLegend | #405721 | 1:200 |
| IgM | PE-Cy7 | II/41 | eBioscience | #25-5790-81 | 1:200 |
| IgM | APC/eFluor780 | II/41 | eBioscience | #47-5790-82 | 1:200 |
| IgG1 | FITC | RMG1-1 | BioLegend | #406606 | 1:200 |
| Igκ | APC-Cy7 | RMK-45 | BioLegend | #409503 | 1:400 |
| Igλ | FITC | R26-46 | BD Biosciences | #553434 | 1:200 |
| FAS/CD95 | BV421 | Jo2 | BD Biosciences | #562633 | 1:800 |
| CD38 | PE-Cy7 | 90 | eBioscience | #25-0381-80 | 1:200 |
| IgD | PE | 11-26c.2a | BioLegend | #405705 | 1:200 |
| Streptavidin | BV421 | N/A | BioLegend | #405226 | 1:200 |
| CD16/32 (FC block) | N/A | 2.4G2 | Bio-X-Cell | #BE0307 | 1:200 |

### Statistical analysis and software

Statistical tests used to compare conditions are indicated in the figure legends. Statistical analysis was carried out using GrahPad Prism v.9. Flow cytometry analysis was carried out using FlowJo v.10 software. Graphs were plotted using Prism v.9, and edited for appearance using Adobe Illustrator CS.

---

**Supplementary Figure 1.**
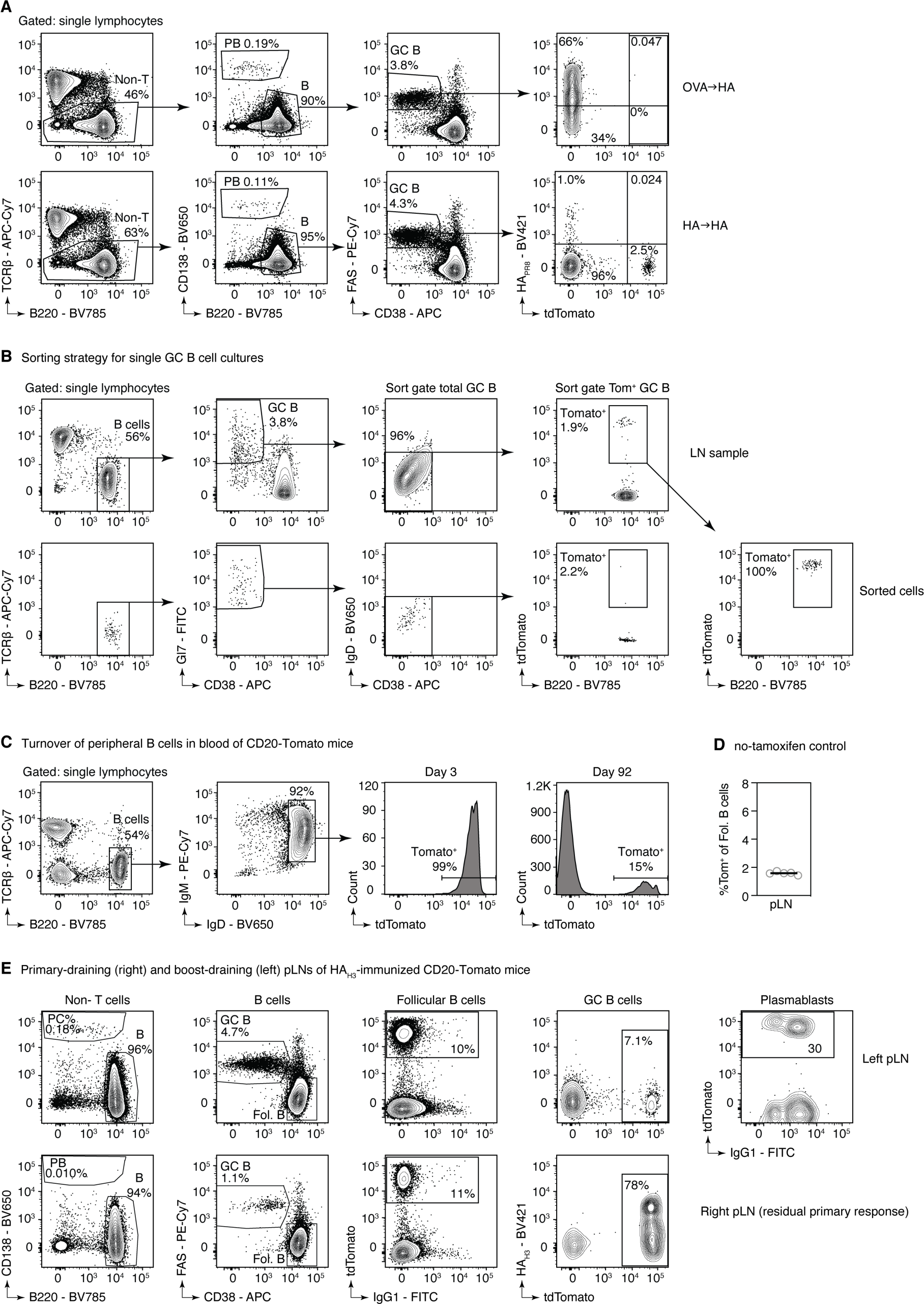
**(A)** Gating strategy for Fig. 1A-F. **(B)** Sorting strategy for single GC B cell cultures, for Fig. 1G-H. Input LN sample (top row) and index sort of sorted cells (bottom row) are shown. B cells were sorted based on GL7^+^CD38^-^IgD^-^ (total GC B) or with an additional gating step on tdTomato (Tom^+^ GC B). **(C)** Gating strategy for naïve B cell turnover shown in Fig. 1I-J. **(D)** Flow cytometry analysis of background tdTomato expression in follicular B cells in pLNs of CD20-Tomato control mice not given tamoxifen. The solid line represents the median. **(E)** Gating strategy for CD20-Tomato recall experiment shown in Fig. 1 K-M. The boosted pLN (top row), and residual primary response in the same mouse (bottom row) are shown.

**Supplementary Figure 2.**
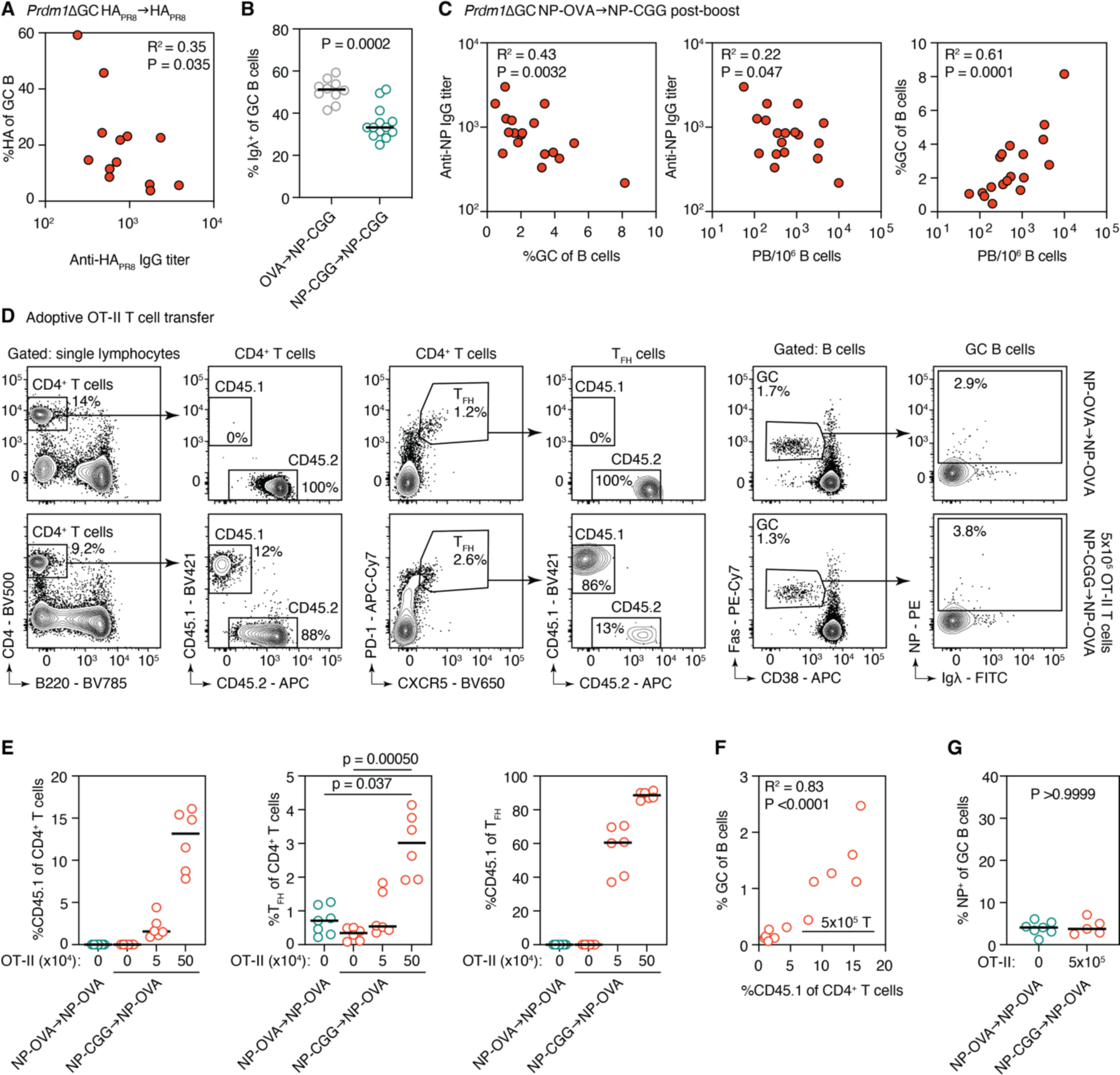
**(A)** Correlation between anti-HA IgG pre-boost titers achieved by depletion and HA-tetramer binding of GC B cells in *Prdm1*ΔGC mice homologously boosted with HAPR8. **(B)** Flow cytometric quantification of Igλ usage by GC B cells. **(C)** Correlation between anti-NP IgG titers and GC formation (left panel) and PB induction (middle panel) in *Prdm1*ΔGC carrier-switch (NP-OVA → NP-CGG) mice, after boost. Correlation between GC and PB formation of same mice (right panel). **(D)** Gating of adoptive OT-II T cell transfer experiment shown in Fig. 4H-I. Homologous recall (NP-OVA → NP-OVA, top row) and carrier switch (NP-CGG → NP-OVA) mice that received 5×10^5^ T cells (bottom row) are shown. Gating of CD4^+^ follicular T helper cells (TFH) and host (C45.2) versus transferred (CD45.1) cells among total CD4^+^ T cells and TFH is shown. B cells were gated as CD4^-^CD8^-^CD138^-^B220^+^, similar to Supplementary Fig. 1A. **(E)** Quantification of the percentage of transferred cells among total CD4^+^ T cells (left) and TFH (right) as well as the quantification of TFH (middle), shown in (D). **(F)** Correlation between the percentage of transferred T cells versus the rescue of GC formation, in carrier-switch mice that received transferred cells. **(G)** Quantification of NP-binding of GC B cells in boost-draining pLNs, shown in right-most panel of (D). The solid line in all bar graphs represents the median. P-values are from Kruskal-Wallis with Dunn’s multiple comparison (E) and Mann-Whitney tests (B, G). R^2^ with corresponding P-values are from Pearson correlation tests (A, C, and F).

